# Mapping Local Conformational Landscapes of Proteins in Solution

**DOI:** 10.1101/273607

**Authors:** M. ElGamacy, M. Riss, H. Zhu, V. Truffault, M. Coles

## Abstract

The ability of proteins to adopt multiple conformational states is essential to their function and elucidating the details of such diversity under physiological conditions has been a major challenge. Here we present a generalized method for mapping protein population landscapes by NMR spectroscopy. Experimental NOESY spectra are directly compared to a set of expectation spectra back-calculated across an arbitrary conformational space. Signal decomposition of the experimental spectrum then directly yields the relative populations of local conformational microstates. In this way, averaged descriptions of conformation can be eliminated. As the method quantitatively compares experimental and expectation spectra, it inherently delivers an R-factor expressing how well structural models explain the input data. We demonstrate that our method extracts sufficient information from a single 3D NOESY experiment to perform initial model building, refinement and validation, thus offering a complete *de novo* structure determination protocol.

## Introduction

In order to function, proteins must adopt a distinct three-dimensional fold. However, a vast range of protein functions, including catalysis, molecular recognition and allosteric signalling, also rely on their ability to adopt various local conformations within this structural scaffold. Understanding these processes therefore requires not only accurate descriptions of protein structures, but also their conformational diversity. NMR spectroscopy is uniquely placed to address these issues, offering atomic-resolution data on samples in native-like physical states. Time averaging of NMR parameters has long been exploited to localise and characterize the timescales of internal dynamics^(1)^. However, the data is also ensemble-averaged over all molecules in the sample volume and should thus provide information on the nature and population of underlying conformational microstates. Accessing this data has long been a goal in NMR spectroscopy^(2-5)^.

Here we aim to elucidate the propensities of individual microstates by means of spectral decomposition. Systematic back-calculation of expectation spectra across a conformational space allows reconstruction of the experimental spectra. In NMR spectroscopy, the richest source of structural data are NOESY spectra, which report on inter-proton distances within a detection limit of 5 to 6 Å. Due to this short spatial range, a large fraction of NOESY intensity can be explained within short, linear sequence fragments. Each such fragment thus represents a sub-space that could be searched systematically to provide detailed information on local dihedral angles and their distributions. Moreover, comparison of the back-calculated and experimental data would provide a quantitative quality measure: an NMR R-factor.

A difficulty in realising this approach lies in the nature of NOESY data itself. The information content of NOESY spectra is very unevenly distributed across the observed intensities, thus small, informative peaks can be overwhelmed by inaccuracies in back-calculation and spectral artefacts. For this reason, quantitative comparison of back-calculated and experimental spectra has been far less applicable in NMR structure determination than equivalent measures used in crystallography^(6-8)^. Here we show that these obstacles can be largely averted in the 3D CNH-NOESY experiment^(9)^. This is an implementation of a ^13^C-HSQC-NOESY-^15^N-HSQC where the indirect proton dimension has been omitted, thus displaying contacts to backbone amide protons in a well-resolved ^13^C dimension. It exploits the higher dispersion and more homogeneous effective linewidths of the heteronuclei, while suppressing water-exchange cross-peaks and obviating the need for stereospecific proton assignments. Crucially, it intrinsically lacks large, uninformative diagonal peaks and the associated baseline and truncation artefacts. Combined, these represent decisive advantages in the accuracy of back-calculation.

In this work we demonstrate that a single 3D CNH-NOESY spectrum contains sufficient information to define population maps of local dihedral sub-spaces. Analytical decomposition expresses the experimental spectra as a linear combination of elements of a features set of back-calculated spectra. In this way, both the reliance on a knowledge-base and the interpretation of spectra in terms of peak or assignment lists can be eliminated. This conformational mapping provides highly detailed data for model building and refinement, with progress monitored by a quantitative R-factor. We validate this method against human Ubiquitin (hUb), widely considered the gold standard for NMR-based protein structure determination^(5, 10-12)^. We further demonstrate the generality of the method by solving the structures of four example proteins.

## Results

### An R-factor from CNH-NOESY data

We have adapted existing routines to back-calculate CNH-NOESY spectra, obtaining 1D ^13^C strips for each backbone amide proton. These are compared directly to equivalent strips extracted from the experimental 3D matrix. An R-factor expressing the discrepancy between experimental and expectation spectra is readily calculated as the fractional root mean squared residual (see Experimental Section). This R-factor is analogous to its crystallographic counterpart, except that it is calculated on a per-residue basis. Our back-calculation routines very accurately reproduce the intensities and line-shapes of experimental CNH-NOESY data collected for hUb (Figure 1a). The back-calculated spectra are also highly sensitive to backbone and sidechain dihedral angles (Figure 1b and Supplementary Figure S1), a prerequisite for conformational mapping.

**Fig. 1.**
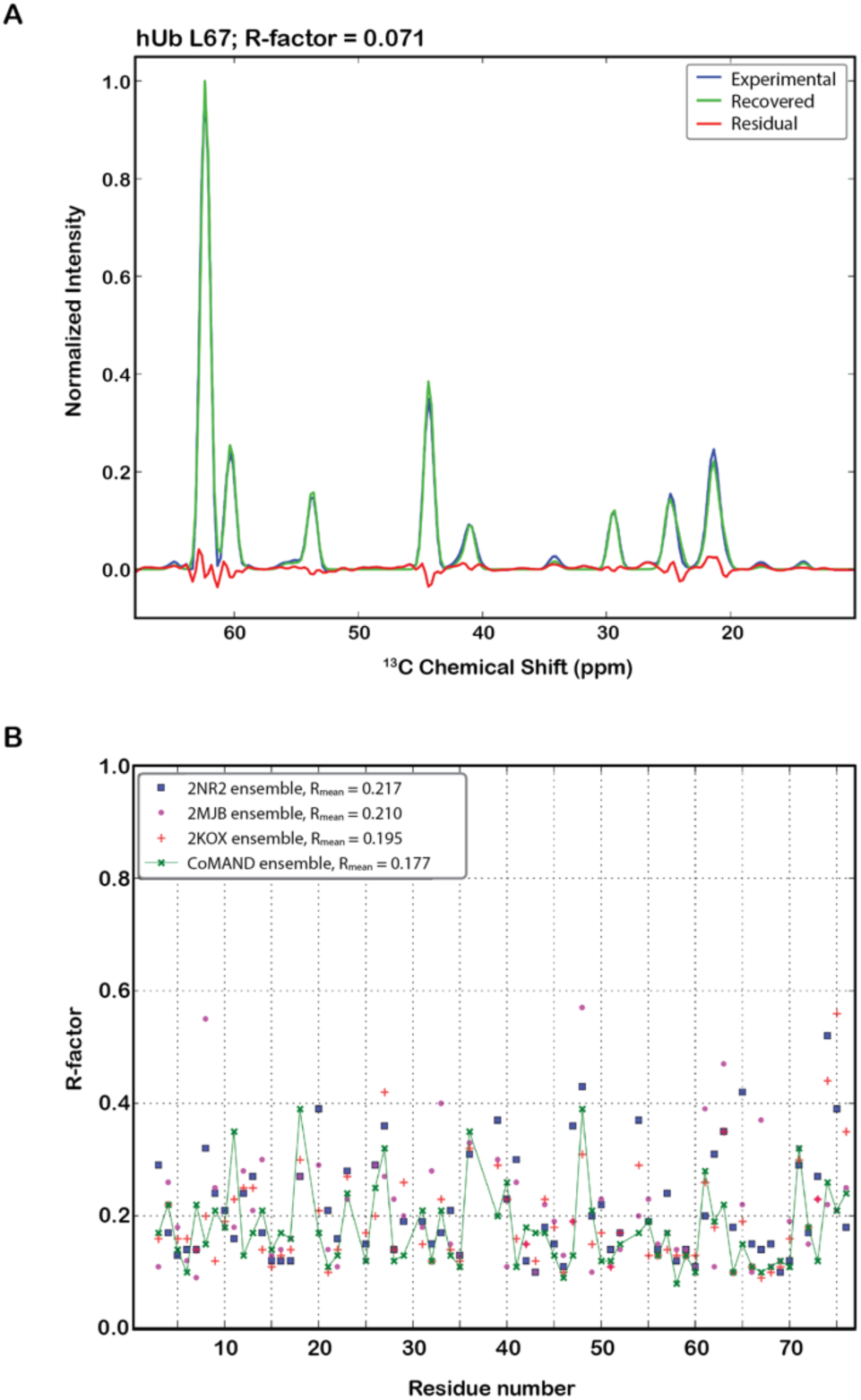
NOESY back-calculation in the CoMAND method. **(A)** An example comparison demonstrating the quality of back-calculation of CNH-NOESY spectra. The experimental spectrum for L67 in human ubiquitin (hUb) is in blue and the recovered spectrum back-calculated by averaging across the 640 models of the 2KOX structure ensemble^(11)^ is in green. The residual signal is in red (R-factor = 0.071). **(B)** R-factors plotted across the sequence for three literature ensembles. These ensembles have been compiled to emphasise different aspects of the hUb structure: 2KOX to elucidate internal motions^(11)^, 2NR2 via a minimal under-restraining, minimal over-restraining procedure^(5)^ and 2MJB to represent a static average pose^(12)^. The average structures for all three ensembles are very similar and differences in R-factors are therefore attributable to the different representations of conformational diversity (see Supplemental Figure S1 for specific examples). These are compared to the CoMAND ensemble (green line).

NOESY intensities are time and ensemble averages over the conformational microstates sampled during the measurement. For this reason, R-factors improve with an accurate and comprehensive description of the ensemble. We demonstrate this for hUb using three reference ensembles that have been compiled according to different metrics. Two have been compiled to elucidate internal motions: the 2KOX ensemble using a large set of residual dipolar coupling (RDC) data^(11)^, and the 2NR2 ensemble according to minimum under-restraining, minimum over-restraining criteria and including 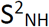 order parameters derived from relaxation data^(5)^. The third, 2MJB, provides a static control ensemble close to the average structure^(12)^. High R-factors are obtained where these ensembles either over- or under-estimate the conformational diversity. Moreover, the use of a residue-wise R-factor in evaluating the ensemble can localize such diversity (Figure 1b).

### Mapping local conformational spaces

For the CNH-NOESY, the vast majority of cross-peak intensity can be explained by intra-residue contacts and those to the immediately preceding residue. The R-factor for residue *i* is thus strongly dependent on conformation in a shifted Ramachandran space defined by the backbone dihedral angles ψ_i-1_ (here denoted υ_i_) and ϕ_i_. This is extended by including the relevant sidechain rotamers up to 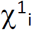 and 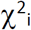, representing a periodic space within a dipeptide fragment that can be searched exhaustively (Figure 2a). The back-calculated spectra for these systematically sampled conformers constitute a features set that can be used to decompose the experimental spectra (Figure 2b). Here we characterize the solution ensemble as a linear combination of elements of the features set, weighted by their respective populations. Calculating these weights is analogous to parts-based representation of complex spectral mixtures often encountered in other fields of spectroscopy^(13)^.

**Fig. 2.**
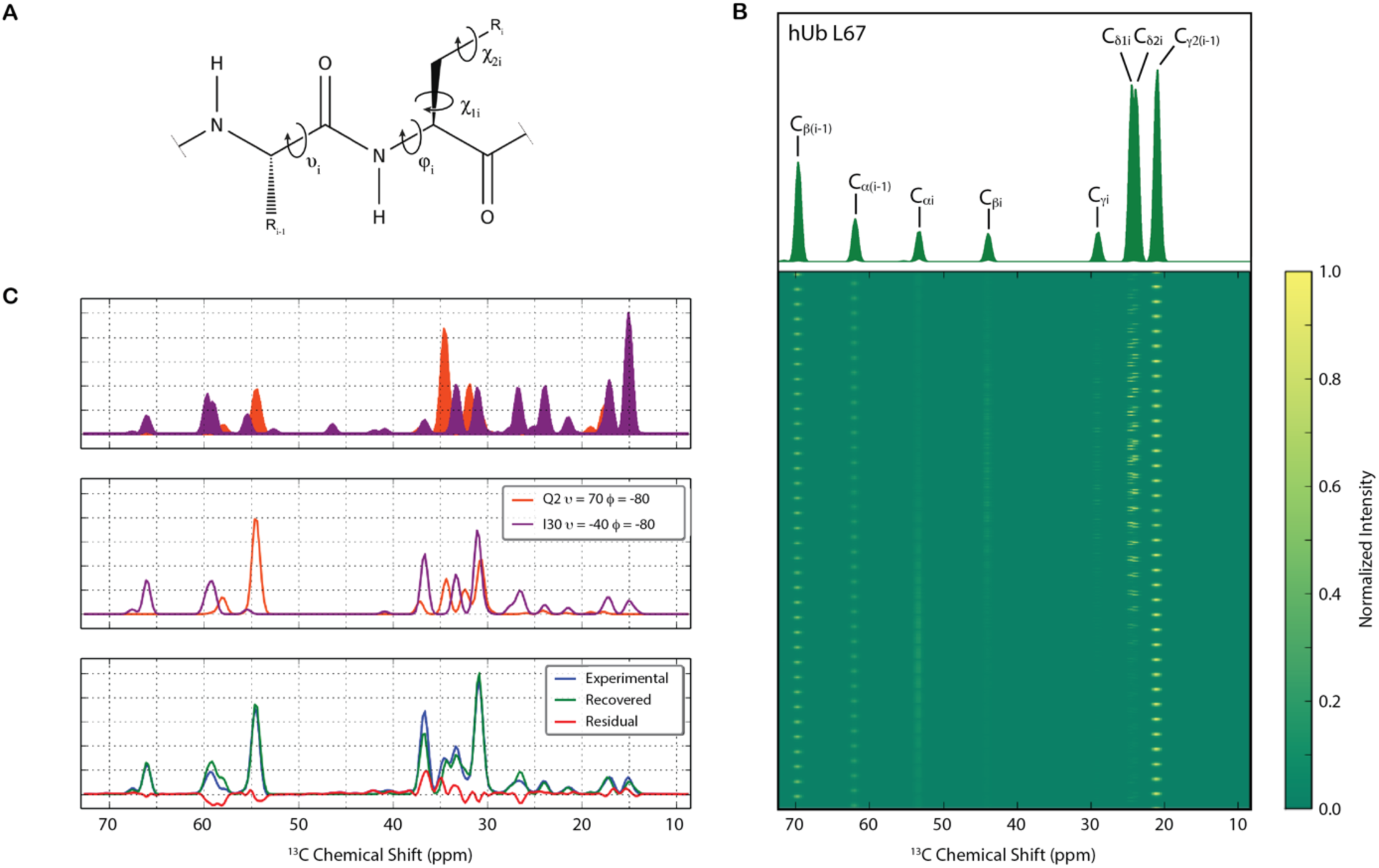
Conformational mapping in the CoMAND method. **(A)** The definition of the conformational search space. Note that in the shifted Ramachandran space, υ_i_ is equivalent to ψ_(i-1)_. **(B)** The features matrix for L67 in hUb shown as a stacked plot where each row is a spectrum back-calculated via systematic conformational sampling, resulting in periodic intensity patterns. The order of sampling, from fastest to slowest, is χ^2^, χ^1^, υ, ϕ with 10° steps for backbone and 120° steps for sidechain angles. The intensity of each peak in the spectrum displays a different dependency on the dihedrals, underlining the power of the data to discriminate individual conformations. The projection of this plot – i.e. all members of the features set overlaid - is shown above with individual peaks assigned. **(C)** Decomposing overlapped spectra. The top panel shows all members of the concatenated features set for Q2 (orange) and I30 (purple) in hUb. Two-component factorization successfully decomposes the completely overlapped experimental spectra, yielding the correct conformations of the respective residues (middle panel).

Decomposition of the experimental spectrum can be framed as a positive matrix factorisation problem^(14)^. The features matrix is represented by ***W*** comprising back-calculated spectra for *l* conformers, resolved to *m* points along the ^13^C dimension. A solution can thus be found for a vector of weights ***H*** in order to reconstruct the observed experimental spectrum ***V***:

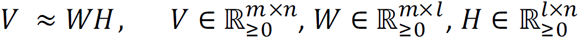
where n=1 if one spectrum is considered per residue. To map the energetic landscape of the conformational space, the yielded estimates of the population weights can in turn be expressed via a Boltzmann factor relative to a reference conformer (Materials and Methods). Examples of these conformational maps for hUb are shown in Figure 3.

**Fig. 3.**
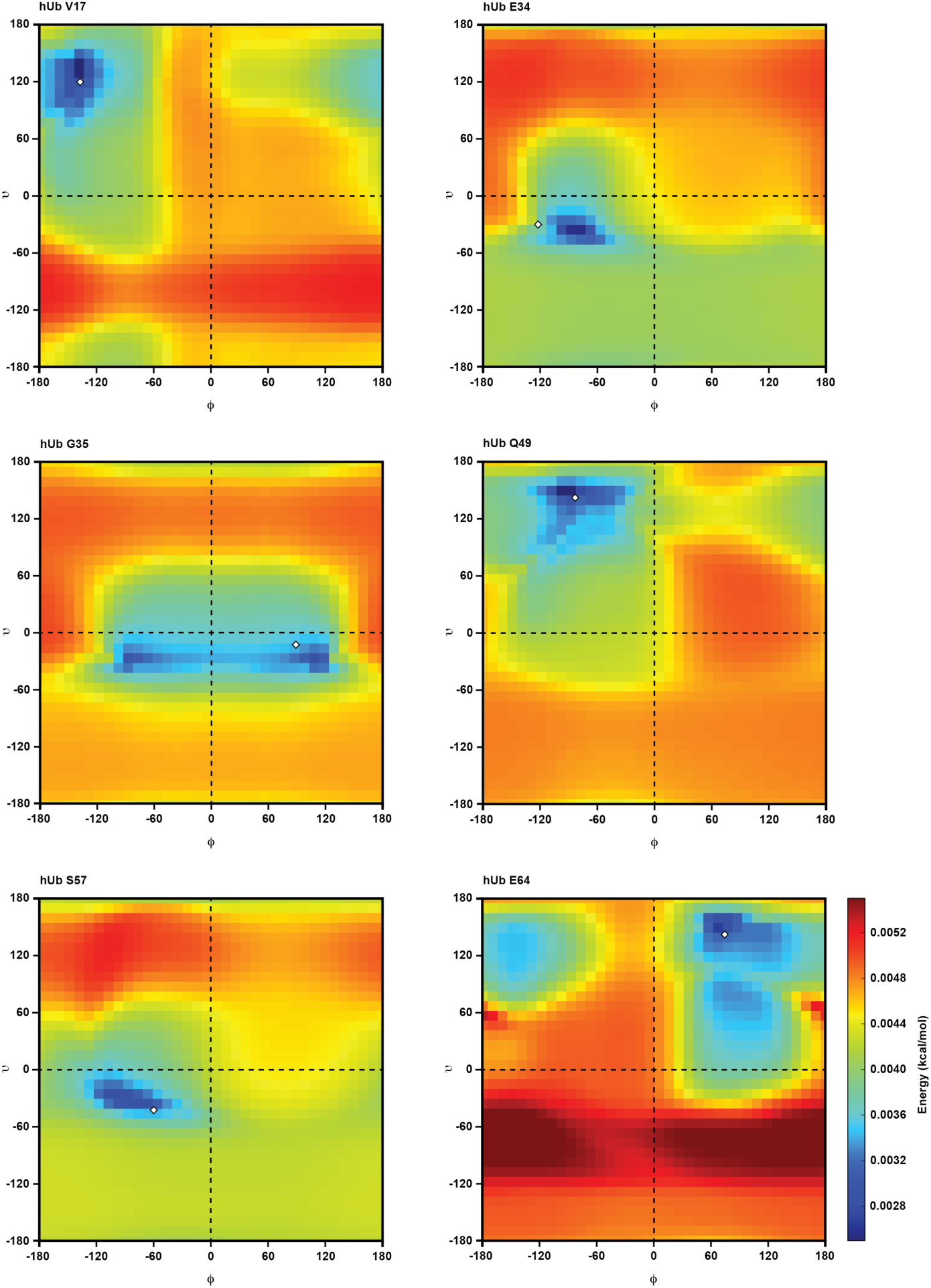
Conformation maps from spectral decomposition. Conformational maps are shown for six examples residues representing different secondary structural contexts in hUb. They are expressed as heat maps of conformer free energy change, relative to a two-conformer global minimum reference state. For non-glycine residues, a two-dimensional (υ, ϕ) slice through the full three- or four-dimensional map at the minimum χ^1^/χ^2^ position is shown. The map for G35 displays typical pseudo-symmetry about the ϕ=0 axis due to the achiral nature of glycine residues. In each map, the minima agree very well with the corresponding crystallographic conformations (1UBQ; white diamonds).

An inherent question in factorization methods is the uniqueness of the solution. Here uniqueness is limited by cross-peaks that cannot be explained within the dipeptide space, but overlap with peak positions in the features set. The larger the fraction of such contamination, the more difficult finding a unique solution becomes. Given the low intensity of potentially contaminating peaks and the high dispersion of the ^13^C dimension, the extent of contamination is usually minor. The extreme case of contamination is the coincidence of ^15^N-HSQC positions for two or more residues, resulting in overlap of the experimental strips. Such a situation can be solved by concatenation of the features sets of the overlapped residues and solving in a multiple-dipeptide space. Figure 2c shows a typical example of this situation, where conformational maps have been obtained for two overlapped residues.

### Structure determination with conformational maps

The conformational maps obtained from spectral decomposition provide rich information for structure determination. At the simplest level, global minima can provide local torsion angles sufficient for model building. These initial dihedrals constitute an agnostic starting point, as they are derived directly from the data without recourse to heuristics or conformational databases. A unique feature of this method is the deployment of R-factors as an objective convergence test that captures both local and long-range contacts. The latter can be isolated by examining the difference between R-factors obtained for a linear peptide fragment and those from the full, folded model. We term this measure the fold factor (F), and it should be negative if the model explains long-range contacts well. Figure 4 shows that the average fold factor (F_mean_) is a sensitive overall measure of correct folding, while the sequence profile can localize misfolded or poorly defined regions. Owing to this independent measure of convergence, any routine can be used to build initial models. Here we employ either a Rosetta-based protocol (Materials and Methods) or a purpose-designed molecular dynamics routine: Simulated Annealing Replica *Seilschaft* (SARS) for initial model building (Materials and Methods).

In order to construct high-resolution ensembles with accurate description of the underlying microstates, the experimental information contained in the conformational maps can be encoded in two possible representations. The first is a temporal equilibrium ensemble derived from molecular dynamics simulations. Here the conformer probability distribution is imposed via a grid-based dihedral energy correction term^(15)^ (CMAP; Figure 3 and Materials and Methods). These override the standard force field CMAPs with bespoke ones on a per-residue basis, providing an experimentally augmented ensemble representation. The other is to aggregate a set of frames from a generalized ensemble using the standard force field, based on an R-factor selection criterion, providing a wider coverage of the phase space.

**Fig. 4.**
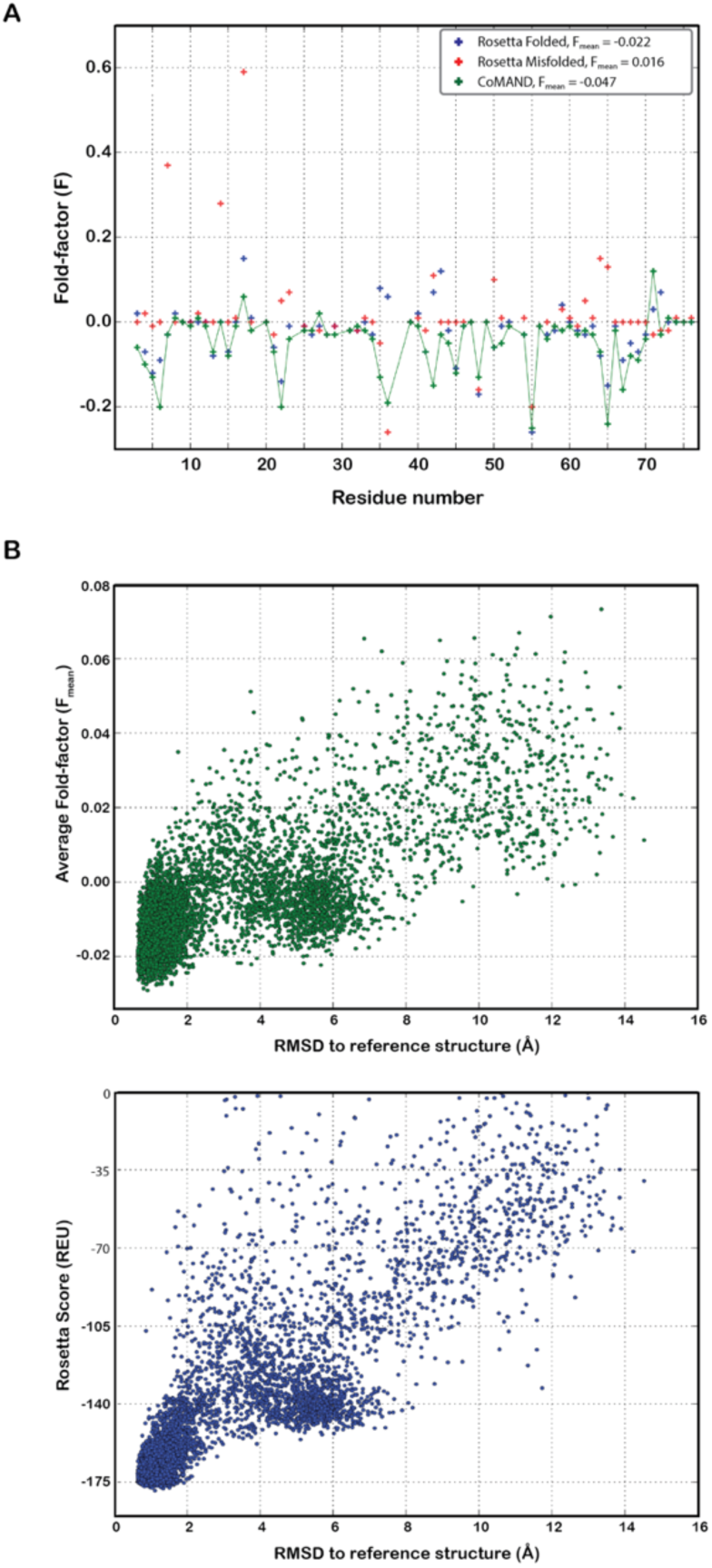
The R-factor as an objective target function. **(A)** Fold factors plotted across the sequence for hUb. Fold factors (F) are calculated on a per-residue basis as the difference between the R-factor calculated for a linear peptide and that calculated for the full structure. This isolates the component of the R-factor not explained by local contacts. Negative values indicate residues in well-folded environments. Values are shown for an initial folded model from Rosetta runs plus a Rosetta structure misfolded by a strand swap in the N-terminal α-hairpin. These are compared to a representative of the final CoMAND ensemble (green line). **(B)** Comparison of the average fold factor (F_mean_) versus the Rosetta score (Rosetta Energy Units) as selection criteria for well-folded hUb models. Both measures are plotted against the RMSD to the reference structure (1UBQ) for the same set of 7215 Rosetta decoys with sub-zero score. Structures with low F_mean_ are consistently close to the reference structure.

We name this method of *de novo* structure determination CoMAND (for <u>C</u>onformational <u>M</u>apping by <u>A</u>nalytical <u>N</u>OESY <u>D</u>ecomposition). In addition to hUb, we present examples for four structure determination projects from our Institute. U3Sfl (125 amino-acids) is a protein designed as a chimera of sub-domain sized fragments, KH-S1 (170 amino-acids) is a fusion construct of the KH and S1 domains of E. coli exosomal polynucleotide phosphorylase. MlbQ is a protein implicated in self-resistance to endogenous lantibiotics in actinomycetes^(16)^. The final example, polb4, is a protein designed to reconstruct the polymerase beta N-terminal domain using two unrelated peptide fragments and is presented here as a *de novo* structure determination.

For all five proteins, we first built starting models. For U3Sfl, KH-S1 and hUb, we extracted backbone dihedral angles from the factorization minima and used these for fragment picking in a Rosetta protocol (Materials and Methods). For MlbQ and polb4 we applied SARS, supplementing the CHARMM36 forcefield with bespoke conformational maps, starting from completely extended chains. For MlbQ, folding was accelerated by the addition of 10 unambiguous NOE distance restraints. We used the average R-factor across the full length of the protein as a criterion for selecting models, choosing a single Rosetta decoy or a single frame from the SARS runs. These models were very similar to respective reference structures (Figure 5). For U3Sfl this was a structure we had previously solved by manual analysis (RMSD over backbone atoms 1.98 Å). For KH-S1, crystal structures are available for homologues of the individual domains (4AM3; 1.48 Å and 4NNG; 1.67 Å). For MlbQ, this was the published solution structure (2MVO; 1.92 Å). As no structure has previously been solved for polb4, we used the design target as a reference structure (RMSD over backbone atoms 1.48 Å).

**Fig. 5.**
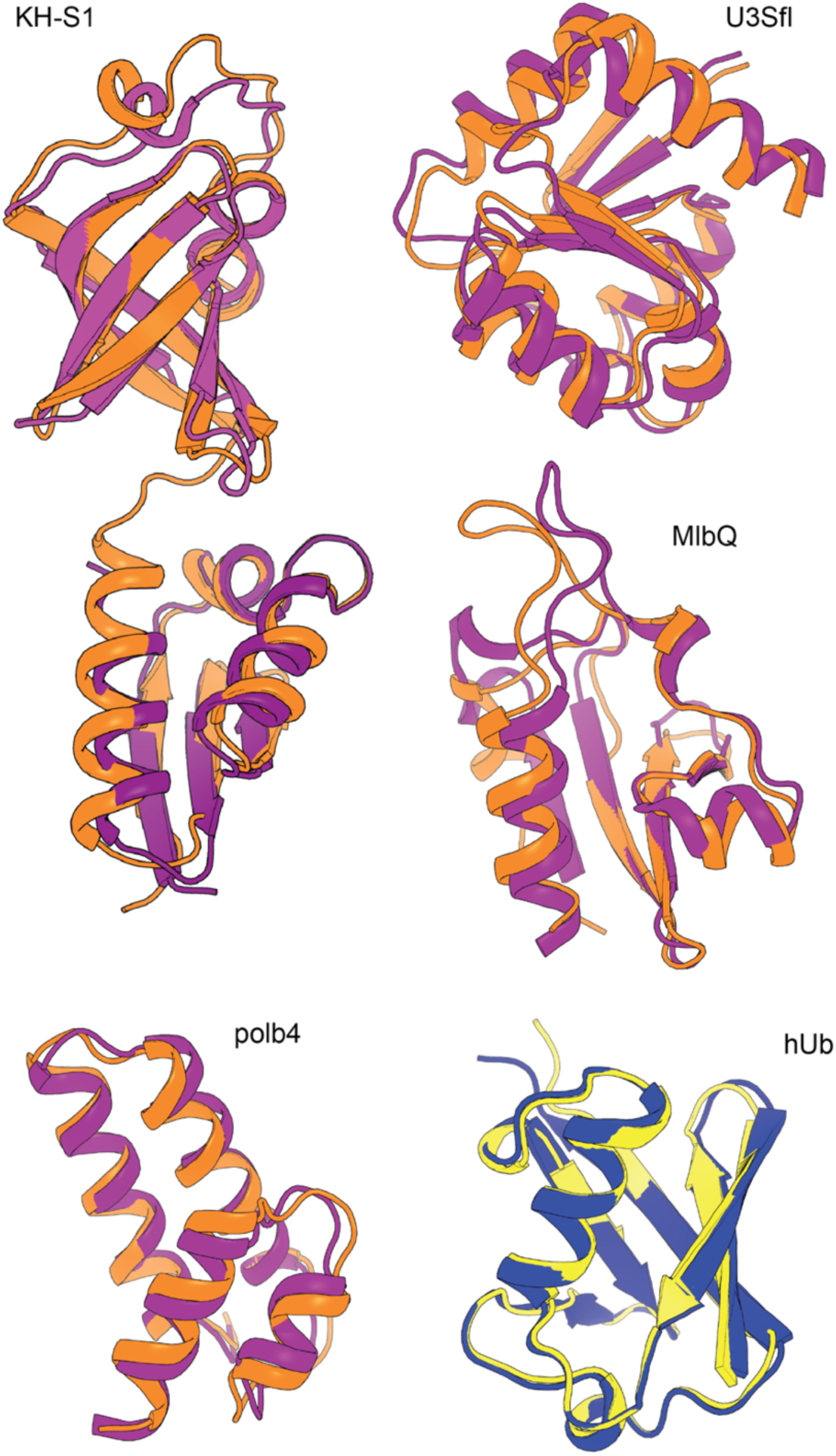
The CoMAND structure gallery. **(A)** The CoMAND structure gallery. Models (orange) are shown superimposed on their respective reference structures. For U3Sfl and MlbQ the reference structures are previously solved solution structures. For KH-S1, the KH domain reference structure is 4AM3 (light purple) and the S1 domain reference structure is 4NNG (dark purple). The *de novo* structure determined for polb4 is compared to the design target. A single model from the refined CoMAND ensemble for hUb is shown in yellow and the reference structure (1UBQ) in blue (RMSD over backbone atoms 0.49 Å).

For refinement we conducted unrestrained molecular dynamics simulations in explicit solvent for microsecond timescales, seeded by the starting model. This was followed by a frame-picking procedure that employs a greedy optimizer to minimize the average R-factor across the ensemble. Given the wealth of structural and dynamics data available for hUb, we compiled such a refined ensemble and compared it to the reference ensembles (2KOX, 2MJB and 2NR2). The resulting ensemble of 20 conformers shows better correlation to experimental NH order parameters than the literature ensembles. It is also comparable in predicting experimental scalar couplings and RDCs to ensembles that have been built on one or more of these observables plus thousands of NOE restraints (Figure 6 and Supplement). This demonstrates the depth of the structural information captured when NOESY spectra are analysed holistically.

**Fig. 6.**
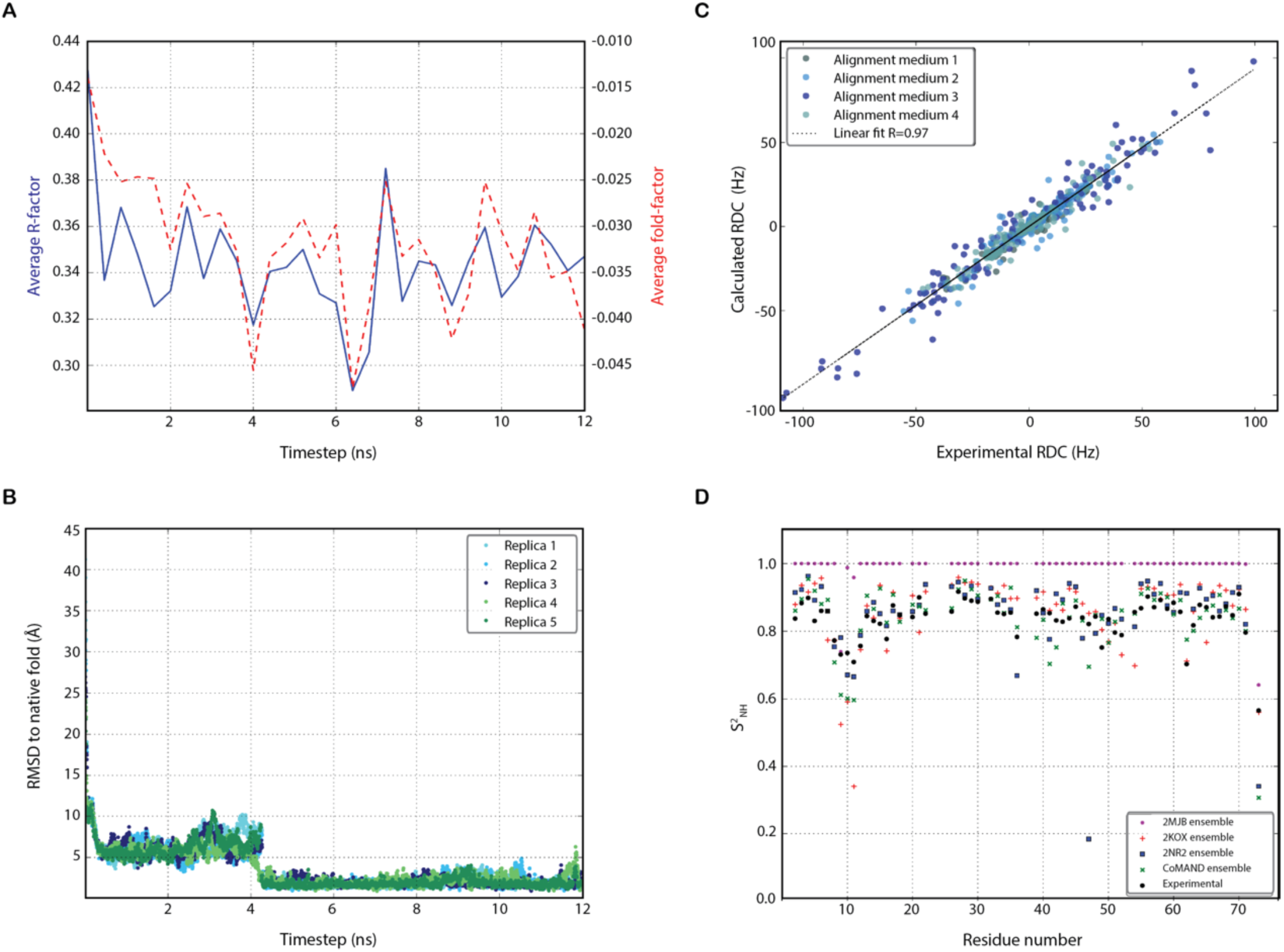
Validation of CoMAND ensembles. **(A)** Evolution of the average sequence R-factor and fold factor along the first replica of the SARS folding trajectory of polb4. The lowest R-factor structure (time point of 6.4 ns) was chosen as a low-resolution model (See also Supplemental Movie S1) **(B)** The RMSD from the native fold along the trajectory of the same SARS folding simulation. Traces for all five replicates are shown, illustrating the convergence of the protocol. **(C)** The CoMAND ensemble independently reproduces NMR observables. The correlation between residual RDC values back-calculated from the CoMAND ensemble and experimental values in four different alignment media is shown^(12)^. The Q-factor expressing the agreement between prediction and experiment for this data set is 0.24. Similarly good agreement is obtained between back-calculated and experiment *^3^J_HNHα_* coupling constants (correlation coefficient = 0.95; Supplementary Figure S2). RDC values report on the orientation of various bond vectors to an external molecular alignment medium and are thus sensitive to both local and global structure. *^3^J_HNHα_* coupling constants report on local ϕ angles. Neither parameter was used in compiling the CoMAND ensemble. **d)** Calculated 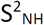 order parameter values across the sequence of hUb using the first 20 models of the 2MJB, 2KOX, 2NR2 and CoMAND ensembles. The CoMAND ensemble best reproduces experimental values derived from NMR relaxation analysis^(29)^ (correlation coefficient=0.82). Correlations for the reference ensembles are shown in Supplementary Figure S2).

## Discussion

NOESY spectra can be seen as an encoding of a proton-proton contact map with an approximate upper distance limit of 5 Å. If correctly decoded as a set of distance restraints, this information is sufficient to solve the structure with high accuracy and precision. However, crowded spectra and the consequent spectral overlap mean that the encoding is ambiguous. Spectral editing, for example via additional frequency dimensions, can only partially alleviate this problem, often at considerable cost in experiment time^(17)^. The consequences of this ambiguity are not only that individual cross-peaks cannot be uniquely assigned – i.e. attributed to a specific proton-proton contact – but also that cross-peaks may comprise significant intensity from several contacts. Conventional NMR structure determination protocols interpret NOESY spectra through peak-picking, assignment and conversion into distance restraints under a paradigm of one peak; one assignment; one restraint. Even automated routines that specifically consider ambiguities will resolve to a single effective restraint per picked peak. This represents a compromise that is not justified by the underlying nature of the data, affecting either the accuracy or precision of distance estimates. In contrast, the CoMAND method makes no interpretation of cross-peaks, and thus stands outside this conventional assignment paradigm.

Given the ambiguity of NOESY data, the incorporation of unambiguous data from other sources is advantageous in NMR structure determination protocols. Particularly useful are data that define local dihedral angles (e.g. scalar couplings), as these are poorly defined by imprecise NOE-based distance estimates. Backbone dihedral angle predictions derived from chemical shift heuristics using the program TALOS are very widely used^(18)^. For example, the CS-Rosetta approach exploits this data to build structural models within the Rosetta framework^(19)^. In contrast to previous methods, we show that direct signal decomposition can yield backbone dihedrals unambiguously and without heuristics. Moreover, we demonstrate that the NOESY data can be leveraged to map the underlying conformational landscape in a systematic fashion. A further key advantage over existing methods is that the whole process of structure determination, including resonance assignment, model building, refinement and conformational mixture elucidation can be objectively assessed by the R-factor as a single metric.

In recent years development of analysis methods in solution NMR of proteins has been driven by the need to make automation more reliable, while using less data and extending the range to larger proteins and more difficult cases, such as membrane proteins. CoMAND contributes to this effort in that it leverages a small set of spectra on a single sample into a high-resolution structure and is therefore applicable where protein concentration or stability are limiting. As the method involves minimum user intervention after the resonance assignment stage, it is also intrinsically suited to automation. However, the most unique feature of the method lies in the power to obtain accurate descriptions of protein conformational ensembles. We therefore anticipate that the method can be applied to studying ligand binding and allosteric processes, promising to elucidate subtle conformational changes in an unprecedented level of detail.

## Materials and Methods

### NMR Spectroscopy

Backbone and sidechain assignments for *de novo* structure determinations were obtained using standard triple resonance experiments. For human Ubiquitin literature values were used. Slight correction of ^13^C shifts against the respective CNH-NOESY spectra was necessary to account for calibration differences between spectrometers and spectrum types. 3D CNH-NOESY spectra were acquired at 800 MHz on a Bruker AvanceIII spectrometer equipped with room temperature probehead. Indirect ^13^C dimensions were typically acquired with ~100 time increments and processed with linear prediction and zero filling to 256 data points. The ^13^C sweep width was set to cover aliphatic carbon resonances; i.e. ~10-73 ppm, resulting in a resolution of ~30 Hz per point. At this resolution, ^1^J_CC_ couplings are unresolved and the spectra were run in non-constant time mode. Broadband ^13^C pulses were used to excite aromatic resonances and these were folded into the aliphatic window without phase inversion.

CNH-NOESY spectra were analysed by extracting one-dimensional ^13^C sub-spectra chosen from a search area centred on assigned ^15^N-HSQC positions (typically 1-3 points in each dimension). As these sub-spectra contain only cross-peaks to a specific amide proton, choosing the strip with highest integral maximises the signal-to-noise. Residues with overlapping search areas were examined separately. In most cases strips with acceptable separation of signals could be obtained. Where this was not possible the residues were flagged as overlapped and a joint strip constructed by summing those at the estimated maxima of the respective components. A set of strips well separated from assigned HSQC positions were averaged to define a global noise level for the spectrum.

### NOESY back-calculation

In order to back-calculate 3D CNH-NOESY spectra we modified the program SPIRIT^(20)^ by porting it to C++ and extending it to accommodate any combination of proton and heteronuclear dimensions. We name this program SHINE, for Simulation of Hetero-Indirect NOESY Experiments. The calculations are based on a full relaxation matrix and thus account for spin diffusion in static structures. Internal motion of the protein is treated by ensemble averaging over *n* contributing microstates, effectively applying an *n*-state jump model of motion, where the life-time of a microstate is assumed to be long compared to the interconversion time. This does not account for true time-averaged phenomena, such as motion-mediated spin-diffusion, which are treated as negligible for the current application. Inputs for the calculations are a chemical shift list, a test structure and a set of simulation parameters. The latter are largely spectral details, such as spectrometer frequencies and sweep widths, which are extracted automatically from the corresponding experimental files (Bruker format), but also include an estimate of a global molecular correlation time. Resonances are modelled as gaussians, with ^13^C linewidths assigned on a class basis, taking into account unresolved ^1^J_CC_ couplings. Here it should be noted that the short acquisition times in an indirect ^13^C dimension (<10 ms on an 800 MHz spectrometer) mean that effective lineshapes are largely governed by apodisation of the time domain. They are thus considerably more homogeneous than for a proton dimension.

The computational demand of NOESY back-calculation depends on the number of protons in the relaxation network. For this reason, we employ sub-structures containing <150 atoms. These can be linear peptides or fragments of a folded structure. Linear peptides are typically tri- or penta-peptides where the test residue is in the second last position. Fragments are compiled at the residue level; residues are included in the substructure if they contain a proton within a given radius (typically 5 Å) of a target residue proton. The output is a one-dimensional strip displaying contacts to a single backbone amide proton. In this mode, back-calculation typically takes less than 20 ms per conformer on a single processor core. The program also outputs a list of peak intensities that can be used to build multi-dimensional spectra suitable for viewing in SPARKY^(21)^, with annotation of individual cross peaks.

Calculation of features sets is performed for each residue for which an experimental spectrum is available. The starting structures are linear peptide fragments extracted from an arbitrary structural model. In the current work these were tripeptides centred on the test residue. For back-calculation of contacts to the amide proton of residue *i* this peptide is modified through a set of torsion angles in a shifted Ramachandran space: angles ψ_i-1_ (here denoted υ_i_) and ϕ_i_ and up to two sidechain χ angles. The backbone angles were searched at 10° granularity, while sidechain angles sample all staggered rotamers. This results in a maximum of 11664 conformers. No checks for steric clashes are applied to test if a conformer is physically reasonable and bond lengths and angles remain constant throughout. An exception is for proline residues where the search space is restricted to a physically realistic range in ϕ_i_. As proline residues lack an amide proton, their features sets are compiled by back-calculation of the amide proton of the previous residue, which is the most sensitive reporter on the υ angle of proline. The set of back-calculated spectra for each residue are stored as a single file. For residues flagged as overlapped, the features set files are concatenated to create a joint set.

### Calculation of R-factors

We define the R-factor as the relative RMS residual between an experimental spectrum and the expectation spectrum back-calculated from a structural model. The use of RMS is in analogy to the quality factor (Q-factor) calculated for residual dipolar coupling data^(22)^. The use of RMS tends to emphasize large outliers relative to the R-factor used in crystallography, which averages absolute differences between experimental and back-calculated structure factors. It also provides a convenient definition of the optimum scaling factor for the back-calculated spectrum *s_cale_*, which can be calculated as:

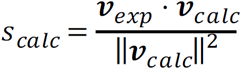
where ***v****_exp_* and ***v****_calc_* are the experimental and back-calculated spectral intensities vectors, respectively. The R-factor is then calculated as:

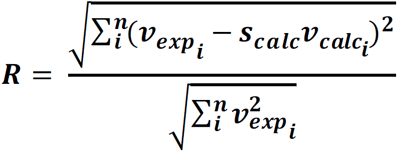

The theoretical range of the metric is from 0 to 1, however the maximum value can only be reached if there is no correspondence between peaks in the experimental and expectation spectra, in which case the scaling factor, *s_calc_*, will approach zero. The practical minimum value is limited by the RMS noise on a per-residue basis.

### Calculation of fold factors

Back-calculation for a structure can be carried out either on a linear peptide or as a fragment of the full structure. The R-factor can therefore be calculated on either basis. Calculation for peptides cannot explain any peaks outside the linear context, whereas all peaks should be explained in a fragment. The difference between fragment and peptide R-factors therefore reports on the fraction of cross-peak intensity that can only be explained in the folded structure. Here we report this difference as a per-residue fold factor, F:

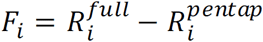
where 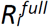 is based on a fragment from the full model and 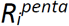 is the penapeptide based R-factor for residue *i*. As R-factors should decrease as more of the cross-peak intensity is explained, the fold factor will be consistently negative for well-folded structures. High positive values are indications of misfolding, while continuous stretches of values close to zero should only be seen for unstructured regions.

### Factorisation and CMAP construction

The dipeptide conformational space was sampled according to the following granularity: Δυ = 10°, Δϕ = 10° Δχ^1^ = 120°, Δχ^2^ = 120°. The experimental vector ***v*** consisted of m = 256 data points of the acquired CNH-NOESY strip (i.e. NOE intensities *vs*. ^13^C chemical shift), while *W* contained all of the back-calculated spectra of the *l* conformers sampled. With the aim of solving for the positive factors vector ***h*** that weights each column of *W* to best explain ***v***. The principal solution can be defined as:

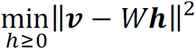
and ***h*** can be also derived directly once the Moore-Penrose pseudo-inverse of the back-calculated spectra matrix, *W^+^*, is computed as:

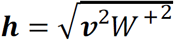

The uniqueness of the solution in positive matrix factorization is limited by the ranks of the component matrices with the upper bound of *min(m,n)*^(23)^. Here, as rank(W) >> rank(***v***) = rank(***v***) = 1, the limiting factor is the rank of ***v***. Thus a 2-conformer block-wise factorisation was sought, being closest to this limit. The 2-conformer solution is also computationally tractable for handling the fine degree of conformational sampling described above. This solution offers a recovered spectrum that has a higher or equal weight against any single-conformer solution. The former two-component weight ***h****_ref_* was used as the highest propensity reference state for estimating the relative normalised propensities of every other available conformer ***h****_i_*. A Boltzmann factor can be directly used to estimate the energy of every conformer according to:

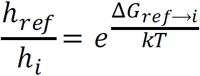

The above procedure was performed across the (υ, ϕ) planes at every (χ^1^, χ^2^) combination. And lowest R-factor yielding plane was the one embedded into the CHARMM36 force field to generate MD ensembles with experimentally derived backbone dihedrals energy surfaces.

### Model building using Rosetta

The Rosetta software package^(24)^ was used to build structural models using backbone dihedral restraints derived from conformational mapping (version 3.6). First, a Rosetta dihedral angle constraint file (.cst) was compiled. For each residue position *i*, a MultiConstraint field was written to comprise both dihedral angles υ_i_ and ϕ_i_. When multiple dihedral angles were possible for one residue position, an AmbigousConstraint field was used to include all possibilities. The Rosetta fragment picking program fragment_picker^(25)^ was used to select 3mer and 9mer fragments satisfying the dihedral restraints from the PDB database. The DihedralConstraintsScore weight used was 500 and the minimum allowed 100. The SecondarySimilarity (weight 150, minimum 1.5) and RamaScore (weight 150, minimum 1.5) both used psipred. FragmentCrmsd was not used. Other fragment_picker parameters were defaults. The fragment database vall.jul19.2011 was searched and 200 3mer and 200 9mer fragments were picked for each position in the protein. The Rosetta ab initio folding program AbinitioRelax was then used to fold the target proteins using these picked fragments (flag file: -ex1, -ex2 -use_input_sc, -flip_HNQ, -no_optH false, -silent_gz 1). Typically, 20,000 decoys were generated for each target protein.

### SARS simulations

The SARS framework was designed to provide robust and efficient conformational sampling without relying on any bioinformatic data, with the only input being CNH-NOESY-acquired dihedral preferences encoded as residue-wise biasing potentials into a standard atomistic force field. The sampling acceleration scheme is based on the assumption that the native structure lies at the global minimum of the potential energy surface that is led to by successively deeper local minima. The algorithm initiates multiple replicas from the same fully extended peptide chain starting system. It then combines alternating rounds of simulated annealing between two temperature baths with conjugate gradient minimisation. Each such round constitutes a search step in a collective swarming behaviour that guides the configuration exchange between replicas whenever a new minimum is reached. In this way, all of the high-energy replicas follow the lead of the lowest energy one. The implementation details and convergence properties of the SARS method will be detailed in a separate publication.

### Molecular dynamics of human ubiquitin

An initial low-resolution model generated by ROSETTA from the CNH-NOESY-acquired dihedral restraints was taken as the input coordinates for molecular dynamics simulations. The standard trajectories were acquired from 10 independent replicas conducted using the standard CHARMM36 force field^(26)^. In contrast, the guided trajectories were acquired from 10 independent replicas where the systems were built using an augmented CHARMM36 with bespoke CMAP potentials. The CMAP potentials were directly constructed on a per-residue basis, in the shifted Ramachandran space (υ_i_, ϕ_i_) from the energy maps as described above. The energy maps where scaled by an arbitrary factor depending on restraint level required, and the standard CMAPs were adopted wherever a bespoke one was not available. These residues with unmodified cross-terms were M1, Q2, G10, P19, E24, I30, P37, P38, G53 and G76.

All of the simulations were performed in explicit TIP3P water as solvent, containing 9226 solvent molecules each, and neutralised by 0.15 M Sodium Chloride in a cubic periodic unit cell. Energy minimisation was performed through 5000 steps of conjugate gradient minimisation, which was followed by 30 ns of NPT equilibration. The time step was set to 2 fs and a Langevin Piston was set to 1 Atm at oscillation period of 200 fs and damping period of 50 fs and a temperature of 298 K. A Langevin thermostat was accordingly set with damping coefficient of 1 ps^−1^. A nonbonded interactions distance cutoff was set to 12.0 Å at a switching distance of 10.0 Å with all nonbonded force and pair list evaluations were performed every timestep, and long-range electrostatics were computed using the Smooth Particle Mesh Ewald method^(27)^ as implemented in the NAMD engine^(28)^. Data was collected from the ensuing NVT trajectories, dumping coordinates every 5 ps for analysis. Frame picking was done from the 10 production trajectories of 30 ns each based on the standard force field that would represent a steady state canonical ensemble.

### Ensemble building

To compile the final CoMAND ensemble for hUb, we performed frame picking from an equilibrium ensemble, such that the compiled conformers belong to microstates of minimal free energy and maximal entropy at the target temperature. This should provide more physically realistic final models compared to those collected from constrained tempering schemes with unrealistic Hamiltonians. To pick frames from the production trajectories we applied a greedy algorithm aimed at minimising the average R-factor for all residues where experimental CNH-NOESY strips were available. For consistency, residues lacking experimental strips (M1, Q2, G10, P19, E24, I30, P37, P38, G53, G76) were excluded from the following comparisons with other datasets.

### Validation versus NMR observables

Expectation NMR observables, R-factors and fold-factors were calculated for the CoMAND and reference hUb ensembles. For uniformity, only the first 20 conformers from each ensemble were considered for the comparisons shown in the figures. For backbone amide order parameters, frames were aligned into a singular molecular frame of reference that best fits backbone atoms of residues 1 through 70. The order parameter, 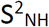, was directly computed according to the method described by Nederveen and Bonvin^(29)^ through the following equation:

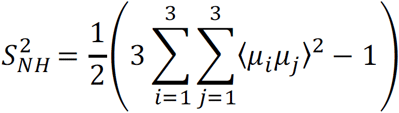
where μ_i_ is the normalised internuclear NH bond vector in the molecular frame of reference and 〈*〉 represents the ensemble-averaged value. Expectation H^N^-H^α^ scalar coupling constants were calculated according to the following Karplus function:

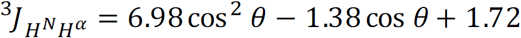
where θ is the H^N^ – N – C^α^ – H^α^ dihedral angle. These were compared to literature experimental values^(30)^. Deviations from experimental residual dipolar couplings and an overall Q-factor for the CoMAND ensemble were calculated using the program PALES^(31)^ using 996 backbone coupling published for 2MJB across four alignment media^(12)^.

## Acknowledgments

We thank Birte Höcker and Remco Sprangers and their co-workers for providing samples for U3Sfl and KH-S1. Programming of the SHINE package was supported by Professors Alois Knoll and Horst Kessler at the Technical University, Munich. We thank Andre Noll (MPI for Developmental Biology, Tübingen) for helpful discussions.

**Author contributions:** MG designed and implemented spectral decomposition routines and the SARS protocol, performed molecular dynamics calculations and analysed their output. MC implemented back-calculation routines. MR programmed SHINE. HZ performed Rosetta calculations and analysis. VT and MC acquired and analysed NMR spectra. MC conceived and directed the project. The manuscript was written by MC and MG and edited by all the authors.

**Declaration of Interests:** This work was supported by institutional funds of the Max Planck Society. The funders had no role in study design, data collection and analysis, decision to publish, or preparation of the manuscript.

**Data availability:** Coordinates for the CoMAND ensemble for human ubiquitin have been deposited in the Protein Data Bank under the accession number (TBC).

## Supplementary Materials

Fig. S1. CNH-NOESY based R-factors are highly sensitive to local conformation.

Fig. S2. The CoMAND ensemble independently reproduces NMR observables.

Fig. S3. Fold-factors identify well-folded models for hUb.

Fig. S4. The CoMAND ensemble for human ubiquitin.

Movie S1. The SARS folding trajectory of polb4.

## Supplementary Materials

**Figure S1:**
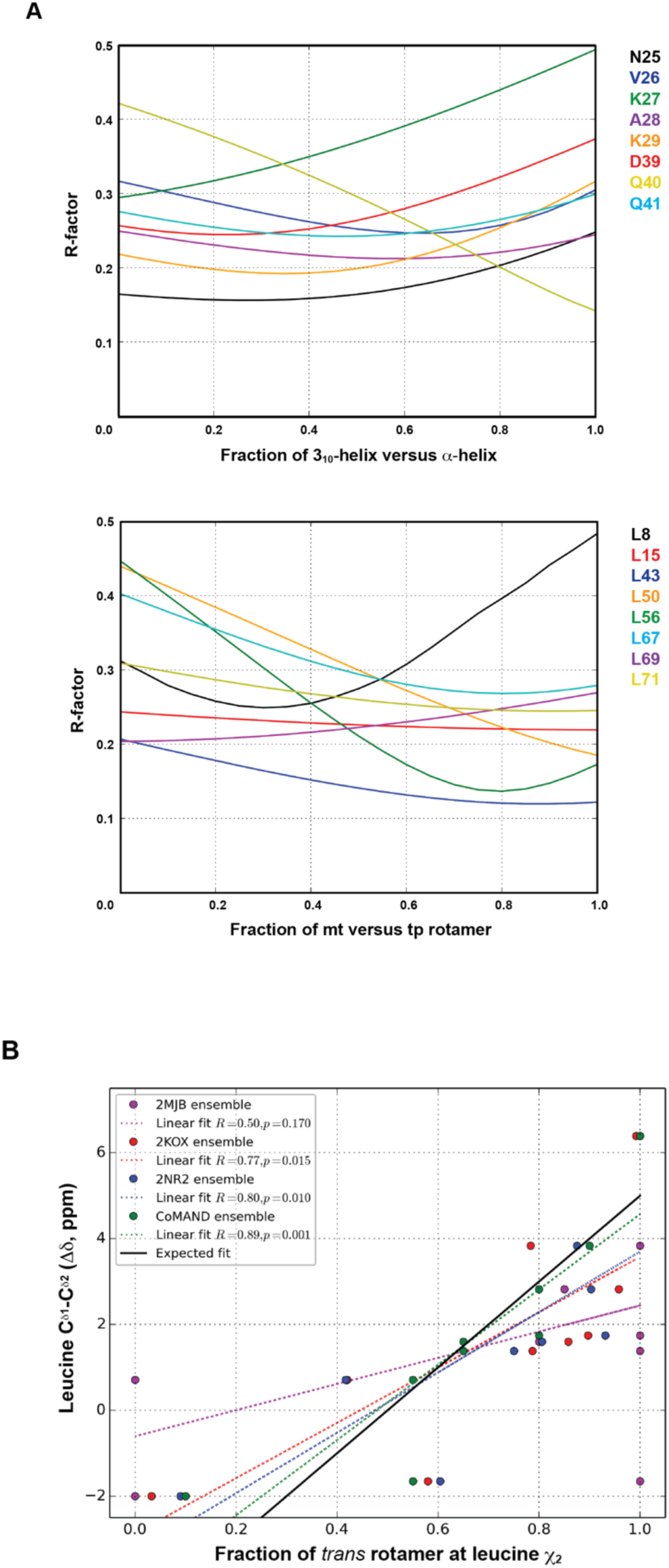
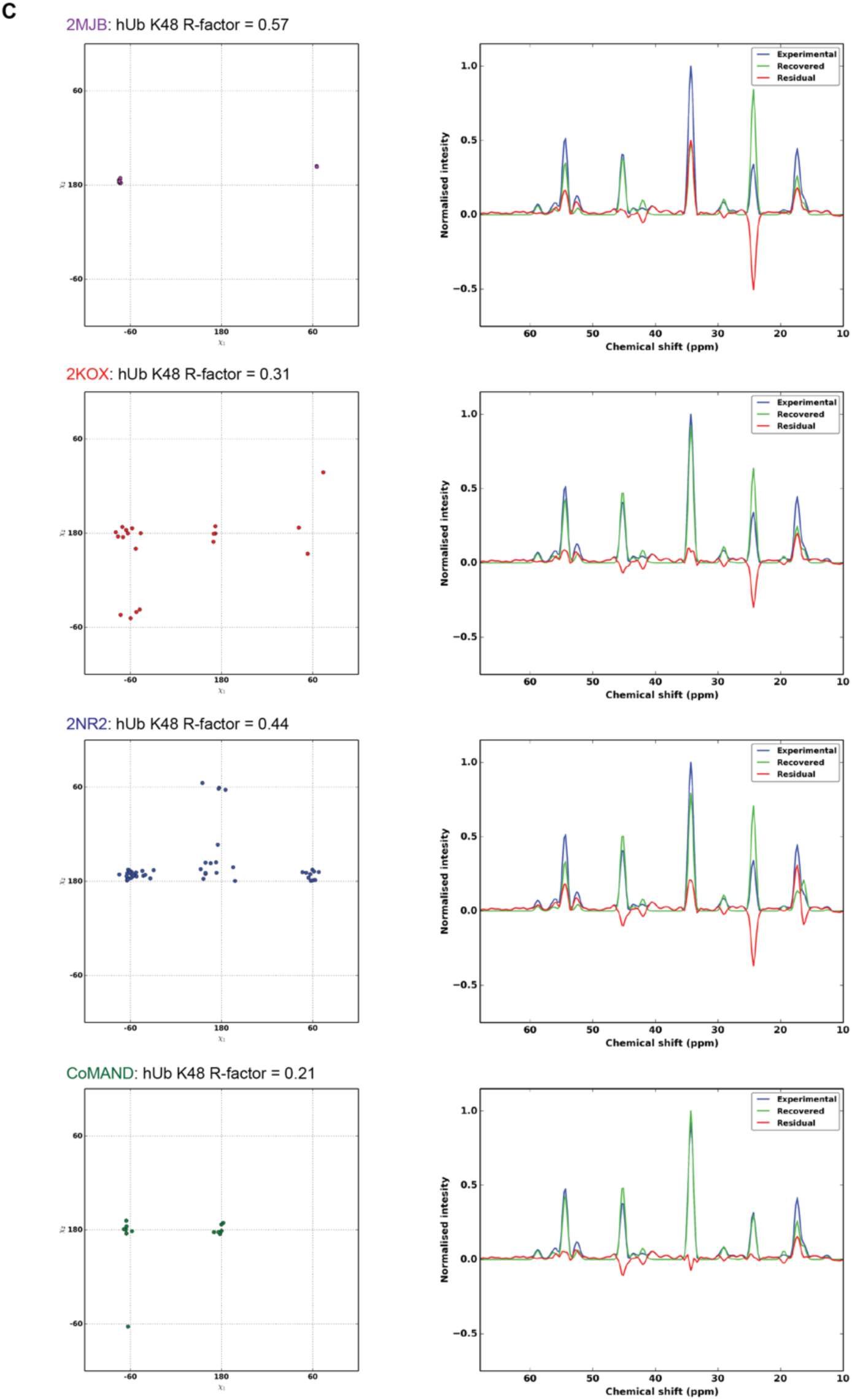
CNH-NOESY based R-factors are highly sensitive to local conformation. **(A)** Plots are shown of R-factors versus composition for idealised two-state conformational mixtures. The upper plot shows linear mixtures between α-helical (υ = −40°, ϕ = −62°) and 3_10_-helical (υ = −10°, ϕ = −95°) conformations, modelling a conformational transition by helical unwinding. Traces are shown for selected residues in helical regions of human Ubiquitin (hUb). Some residues are best explained by pure conformers, e.g. K27 and Q40, while others have R-factor minima for mixtures. The lower panel models transition between the two most populated sidechain rotamers for leucine residues: mt (χ^1^ = −60°, χ^2^ = 180°) and tp (χ^1^ = 180°, χ^2^ = 60°). Traces are shown for all leucine residues in hUb for which data is available. Residues sampling multiple conformations are clearly identified. **(B)** Validating conformational mixtures in the CoMAND ensemble. The chemical shifts of leucine C^δ1^ and C^δ2^ carbons are sensitive to the χ^2^ rotamer due to a “γ-gauche effect”^(32)^. The difference in these shifts correlates with the proportion of *trans* rotamer in leucine residues and thus provides an independent estimate of the conformational mixtures described in the lower plot of panel A. The plots show the shift differences (Δδ) versus the proportion of *trans* rotamer for the CoMAND and three reference ensembles for hUb. These ensembles have been compiled according to different metrics: 2KOX to elucidate internal motions, 2NR2 according to minimum under-restraining, minimum over-restraining criteria and 2MJB, which represents a static structure close to the average structure and is therefore not expected to explain conformational diversity well. The CoMAND ensemble best explains the observed chemical shifts. The expected shift difference (solid line) is based on the equation derived by Mulder^(32)^. **(C)**. Literature ensembles for hUb over- and under-estimate conformational diversity. The panels on the left show the distribution of χ^1^/χ^2^ rotamers for K48 in hUb in the CoMAND and reference ensembles. For 2KOX only the first 20 models of the ensemble are shown for clarity. The panels on the right show the comparison between experimental spectra and spectra back-calculated over the whole ensemble. The R-factors for these comparisons demonstrate its sensitivity to accurate representation of conformational diversity.

**Figure S2:**
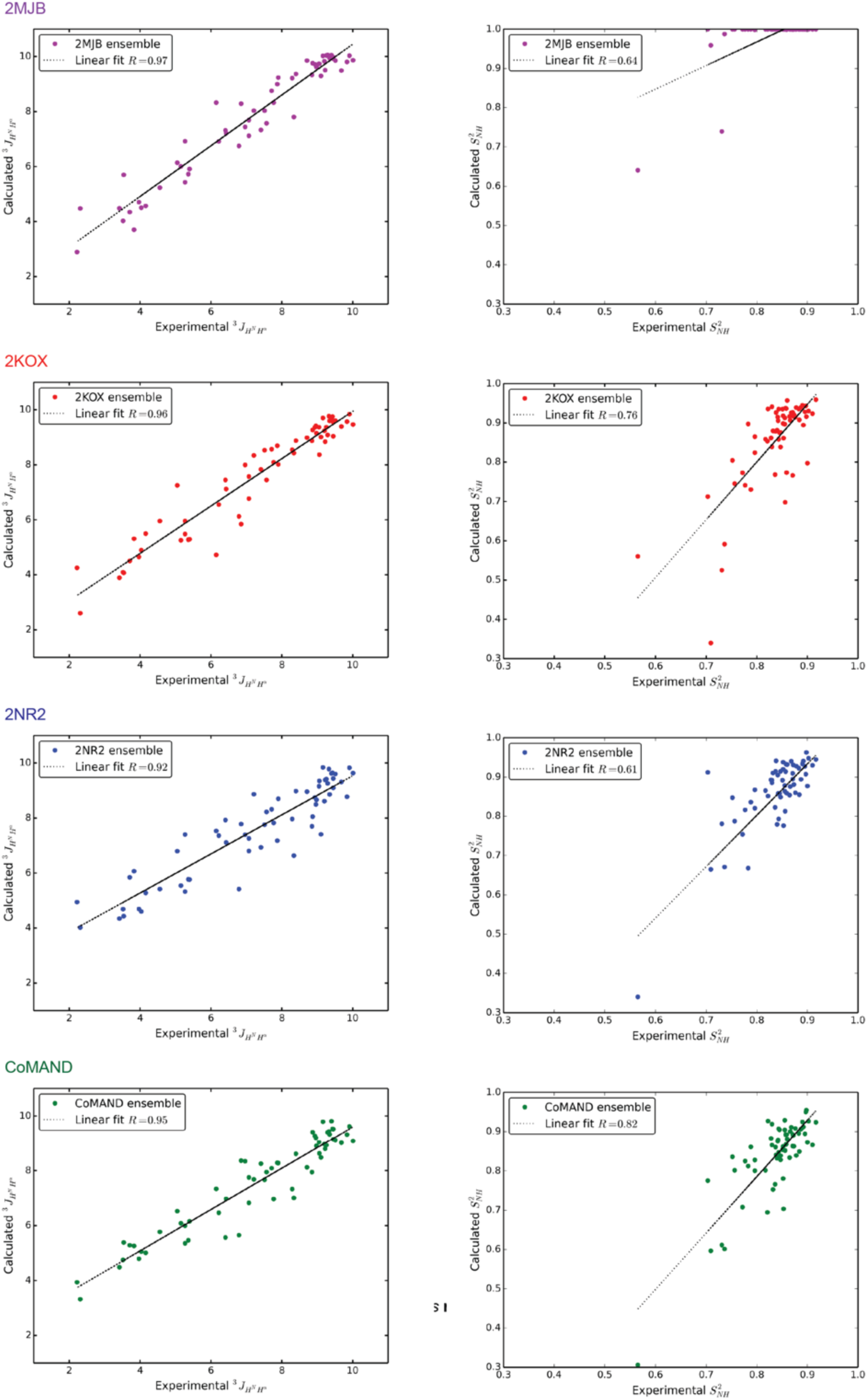
The CoMAND ensemble independently reproduces NMR observables. Correlations are shown between experimental and back-calculated *^3^J_HNHα_* coupling constants^(12)^ and backbone 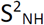 order parameters^(29)^ for the CoMAND and reference ensembles. Note that the 2MJB ensemble was refined against *^3^J_HNHα_* couplings, while 2NR2 was refined against order parameters.

**Figure S3:**
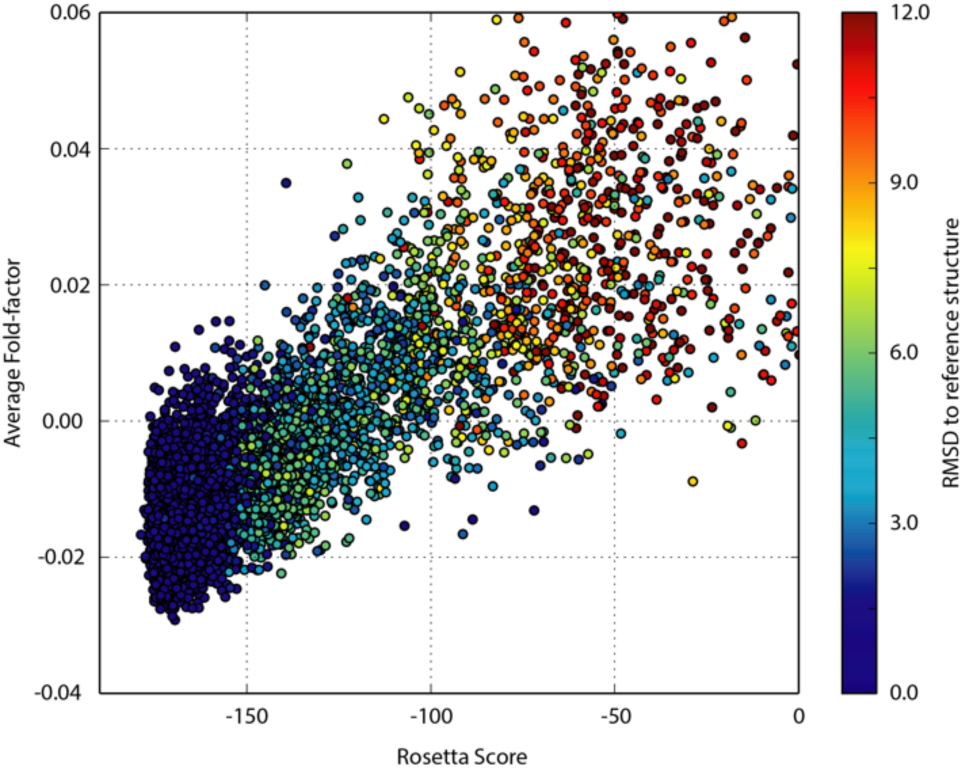
Fold-factors identify well-folded models for hUb. Correlations are shown between the average fold-factor (F_mean_) and Rosetta Score (Rosetta Energy Units) for a set of 7215 decoys with sub-zero score calculated for hUb. Each point is coloured according to the RMSD to the reference crystal structure (1UBQ). Agreement between the two measures is a very good predictor of well-folded decoys.

**Figure S4:**
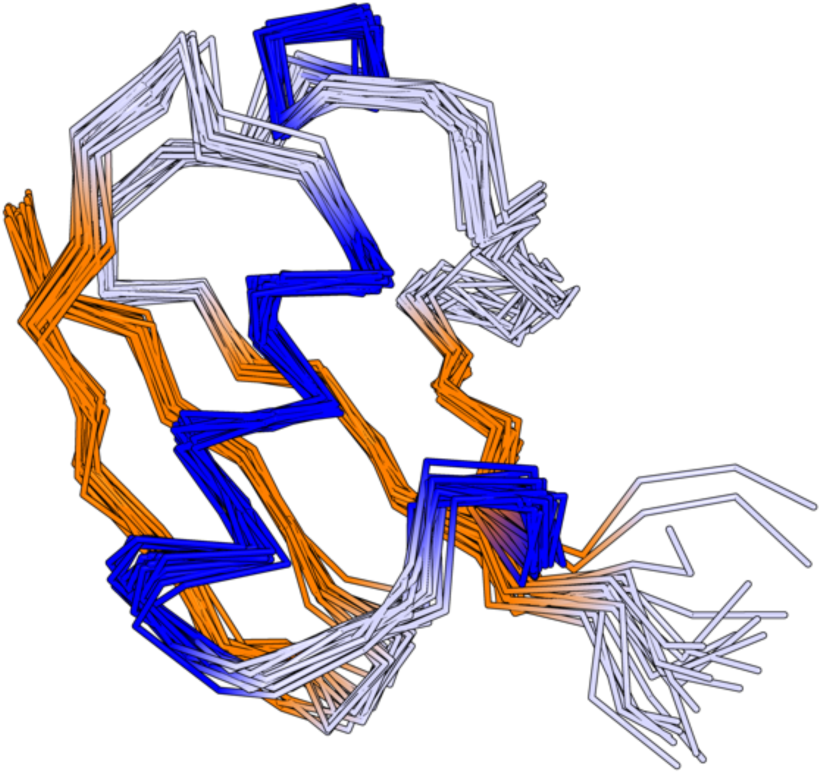
The CoMAND ensemble for human ubiquitin. The refined ensemble for hUb (20 models) is shown superimposed over backbone atoms. Helices are in blue and β-strands in orange. The backbone RMSD to the average structure for the ensemble is 0.64 Å. The ensemble has been compiled by frame-picking structures from an unrestrained molecular dynamics simulation employing a greedy algorithm to minimise the overall average R-factor.

**Movie S1.** The SARS folding trajectory of polb4. The movie shows the time evolution of the communicating replica performing a *seilschaft* search for lower energy minima. The inset shows the backbone RMSD from the design as a function of time.

